# ConvNeXt-Driven Detection of Alzheimer’s Disease: A Benchmark Study on Expert-Annotated AlzaSet MRI Dataset Across Anatomical Planes

**DOI:** 10.1101/2025.07.10.664260

**Authors:** Mahdiyeh Basereh, Matthew Abikenari, Sina Sadeghzadeh, Trae Dunn, René Freichel, Prabha Siddarth, Dara Ghahremani, Helen Lavretsky, Vivek P. Buch

**Author notes:** **Corresponding Author:** Vivek P. Buch, M.D. Department of Neurosurgery, Stanford University School of Medicine Stanford, CA 94305. Indicates co-first authorship (Mahdiyeh Basereh, Matthew Abikenari). **Ethics Statement:** Approval from an institutional review board (IRB) was not required for this investigation as it was carried out on a fully de-identified dataset (AlzaSet) of anonymized structural MRI scans. No personal identifiable data, such as names, facial features, or medical record numbers, was retained. The data set was created from available clinical imaging data that had already been de-identified before analysis, and there was no direct contact with human subjects in processing or data collection. According to institutional and national guidelines, research conducted using such de-identified data is exempt from IRB review. **Data Availability Statement:** The AlzaSet dataset used in this study is available in Zenodo under embargo at the following DOI: https://doi.org/10.5281/zenodo.15564968 The dataset may be obtained by researchers on reasonable request from the corresponding authors. The data consist of anonymized T1-weighted structural MRI scans from Alzheimer’s disease and cognitively normal subjects, carefully curated and labeled by expert neuroradiologists. All releases of the dataset are cited under the above DOI, which will link to the most recent version.

## Abstract

Alzheimer’s disease (AD) is a leading worldwide cause of cognitive impairment, necessitating accurate, inexpensive diagnostic tools to enable early recognition. In this study, we present a robust deep learning approach for AD classification based on structural MRI scans, ConvNeXt, an emergent convolutional architecture inspired by vision transformers,. We further present AlzaSet, a novel, expertly labeled clinical dataset with axial, coronal, and sagittal perspectives from AD and cognitively normal control subjects. Three ConvNeXt sizes (Tiny, Small, Base) were compared and benchmarked against existing state-of-the-art CNN models (VGG16, VGG19, InceptionV3, DenseNet1 21). ConvNeXt-Base consistently outperformed the other models on coronal slices with accuracy of 98.37% and an AUC of 0.992. Coronal views were determined to be most diagnostically informative, with emphasis on visualization of the medial temporal lobe. Moreover, comparison with recent ensemble-based techniques showed that superior performance with comparable computational efficiency. These results indicate that ConvNeXt-capable models applied to clinically curated data sets have strong potential to provide scalable, real-time AD screening in diverse settings, including both high-resource and resource-constrained settings.

## 1. Introduction

Dementia is a rapidly growing global health concern, affecting an estimated 57 million individuals worldwide, with approximately 10 million new cases reported each year. As the seventh leading cause of mortality across all age groups, it represents a significant and rising public health burden [1]. Alzheimer’s disease (AD) accounts for 60–70% of all dementias and is associated with significant burden for both patients and caregivers, as well as the health care systems, and society in general. Addressing AD has become an imperative given the increase in individuals aged over 60 [2]. This demographic pattern calls for preventive healthcare planning and the creation of faster and more accurate diagnostic tools, deployable at scale [3].

AD is a chronic, progressive neurodegenerative disorder marked by structural and functional impairment of the brain, predominantly in the hippocampus and associated cortical regions. The hippocampus, which is a critical structure for learning and memory, is often among the first and most severely affected regions in AD [4]. Hippocampal atrophy, which can be identified by structural magnetic resonance imaging (MRI), is highly correlated with the severity of disease and is an acceptable imaging biomarker [5].

Furthermore, emerging studies increasingly underscore the diagnostic value of peripheral biomarkers when used in tandem with neuroimaging for early Alzheimer’s disease detection. Elevated levels of plasma Aβ40 and Aβ42 have been associated with heightened subjective awareness of memory in older adults with cardiovascular risk, and this has led to the possibility of peripheral amyloid burden being linked with early awareness of cognition [6]. Parallel advances in other neurological domains, such as neuropathic pain, have revealed that integrating deep learning-based imaging and molecular signatures facilitates mechanistic subtyping and stratified treatment in an individualized context [7]. In addition, immune marker remodeling—particularly cytokines and growth factors—has been linked to clinical remission in late-life depression and reflects the importance of inflammatory pathways in neuropsychiatric and neurodegenerative disease courses [8]. Collectively, these findings foster a multimodal diagnostic strategy that integrates imaging, peripheral biomarkers, and computational modeling to increase accuracy and clinical utility in AD diagnosis. Incorporating these molecular pathways alongside imaging biomarkers could enhance prognostic accuracy and therapeutic matching in AD care.

Over the past decade, deep learning (DL) has revolutionized medical image analysis through the delivery of automated, scalable, and extremely accurate solutions to complicated classification tasks. DL is a subcategory of machine learning that utilizes multi-layered artificial neural networks that can learn and derive hierarchical features from data, often without manual preparation of input. In computer vision and medical imaging use, DL-based models have consistently outperformed traditional rule-based and shallow learning approaches [9]. However, one of the biggest constraints in applying DL to medical use is the lack of annotated data due to privacy concerns and resources needed for expert labeling. Transfer learning is an effective method to address the lack of annotated data by leveraging pre-trained models trained on large datasets such as ImageNet [10].

Convolutional Neural Networks (CNNs), the most commonly used DL models used for image processing, represent images in a hierarchical manner with increase abstraction through their hierarcy.. Early layers extract low-level features like edges, gradients, and textures while deeper layers capture higher-order semantic features of objects. Although pre-trained on natural images, CNNs used for image processing demonstrate phenomenal cross-domain adaptability; their hierarchies can readily be fine-tuned to medical imaging domains, hence enabling faster convergence and reducing overfitting on small datasets [11,12].

Contemporary advances in CNN structures like AlexNet, VGG, ResNet, and Inception have attained remarkable performance on many visual recognition tasks. Most recently, ConvNeXt, a model created by researchers at Facebook AI Research (FAIR) in 2022, set new records for convolutional network performance. ConvNeXt is an enhanced ConvNet architecture inspired by Vision Transformer (ViT) design principles. By combining architectural innovation and updated training methods, ConvNeXt achieves best-in-class performance on a variety of computer vision tasks, surpassing performance of both transformer-based and CNN models with computational efficiency [13].

In this paper, we present a ConvNeXt-inspired design for AD detection from structural MRI scans. To encourage and enable reproducibility in the field, we introduce AlzaSet, a new expert-annotated dataset of AD patients and healthy, cognitively normal controls. Using this approach, we aimed to provide more accurate and accessible diagnostic support tools for clinicians and radiologists, with a powerful tool in hand to diagnose AD at an earlier stage and reduce rates of underdiagnosis in the clinic.

## 2.0 Background and Prior Work

Deep convolutional neural networks (CNNs) are now normative in medical image analysis, particularly in neuroimaging applications such as Alzheimer’s disease (AD) detection. Their hierarchical structure entails automatic feature learning from imaging data, with deeper layers containing more abstract representations. Transfer learning from large-scale pretrained models has in recent years become a feasible solution for addressing the limitations due to small medical datasets and making successful model generalization with limited data possible.

CNNs originally developed for classifying natural images have been found to be successfully adapted for neuroimaging with transfer learning. Liu et al. [14] employed a pre-trained model of AlexNet to differentiate MR images into pathological or non-pathological and reported 100% accuracy using a Harvard Medical School dataset (N=?). Following this, Khan et al. [15] optimized VGG19 using MRI data from the Alzheimer’s Disease Neuroimaging Initiative (ADNI), achieving 99% accuracy by employing a combination of entropy-based image selection and better performance on a small dataset.

Other researchers have implemented comparative assessment across a range of pretrained architectures. A comparison of AlexNet, GoogleNet, and ResNet101 on classification of ADNI data indicated that both AlexNet and GoogleNet achieved area under the curve (AUC) results greater than 89%, and demonstrated clear superiority over CNNs without transfer learning or pretraining [16]. Mehmood et al. [17] obtained 98.73% classification accuracy by employing a variant of VGG19 by freezing different convolutional blocks and using data augmentation to address class imbalance. Similarly, in [18], the authors utilized a stepwise fine-tuning and freezing method with VGG16 and VGG19, further improving performance with manually pre-designed fully connected layers for dementia stage classification. Their models obtained 96.39% and 96.81% accuracy, respectively, and outperformed DenseNet and ResNet50 for the same task.

Ensemble approaches have also been explored to enhance predictive capacity. For instance, [19] proposed two ensemble architectures: one combining VGG19 and Xception (achieving 93% accuracy), and another that combined ResNet50 and VGG16 (achieving 95%). In [20], various models (VGG16, VGG19, AlexNet) were compared with baseline machine learning classifiers support vector machines (SVM) and random forest (RF). The deep learning models, particularly those that used transfer learning, significantly outperformed traditional methods, with accuracies of 98–100% as opposed to 81.6% (SVM) and 85% (RF).

Later works concentrated on employing more recent CNN model architectures such as ConvNeXt for the detection of AD. ConvNeXt was integrated with a pipeline of 3D convolution and 3D Squeeze-and-Excitation (SE) attention in [21] and attained an accuracy of 94.23% for AD vs. normal controls. Another study [22] employed ConvNeXt on the ADNI database to make predictions about AD stages and compared its performance against ResNet50 and Swin Transformer—a high-performance ConvNeXt outperformed the two alternatives with 95.3% accuracy compared to 93.8% (ResNet50) and 94.3% (Swin Transformer), which shows that novel CNN architectures can achieve superior or equivalent performance to transformers in medical image classification tasks with limited training data.

Collectively, these studies demonstrate the ability of transfer learning and modern CNN architectures to predict AD based on neuroimaging. However, the majority of such studies rely on high-dimensional 3D imaging or complex ensemble pipelines that are usually computationally intensive and less convenient for implementation in the clinical setting. On the other hand, 2D slice-based models, particularly those learned across multiple anatomical planes, offer significant advantages in computational cost, scalability, and compatibility with existing imaging infrastructures. While they are straightforward, fewer efforts have comprehensively optimized 2D models like ConvNeXt with domain-specific preprocessing and benchmarking performance along the orientation axis. Our method addresses this gap by employing a customized ConvNeXt-based pipeline on an expert-annotated novel dataset with a state-of-the-art performance and, at the same time, being deployable in real time in high- esource and low-resource clinical settings.

## 3.0 Materials and Methods

### 3.1 Dataset Description: The AlzaSet Cohort

This study introduces a novel, expertly annotated dataset termed AlzaSet, comprising whole-brain structural T1-weighted MRI scans from individuals diagnosed with Alzheimer’s disease (AD) and cognitively normal control (NC) subjects. The dataset includes T1-weighted MRI scans from a total of 79 participants—63 with clinically diagnosed AD and 16 cognitively normal controls. Data collection was conducted in collaboration with the Radiology Department at Sasan Hospital (Tehran, Iran), and all images were reviewed and labeled by experienced neuroradiologists. Imaging was performed using a Siemens MAGNETOM Aera 1.5 Tesla MRI scanner (Siemens Healthineers, Erlangen, Germany), equipped with Tim® 4G integrated coil technology and Dot™ (Day optimizing throughput) workflow protocols. Each MRI session included standard neuroanatomical orientations—axial, coronal, and sagittal planes—to ensure comprehensive spatial coverage of the brain. Demographic information of the participants are presented in Table 1.

**Table 1.** Demographic information of AlzaSet.

| Participants Status | Age | Gender(F/M)% |
| --- | --- | --- |
| AD | $70.1 \pm 11$ | 37/63 |
| NC | $57.7 \pm 18$ | 54/46 |

To facilitate image processing, each frame of the scanned images was converted from DICOM (.dcm) to JPEG (.jpg) format while preserving original dimensions and resolution. See Table 2 for complete numerical distributions.

**Table 2.**
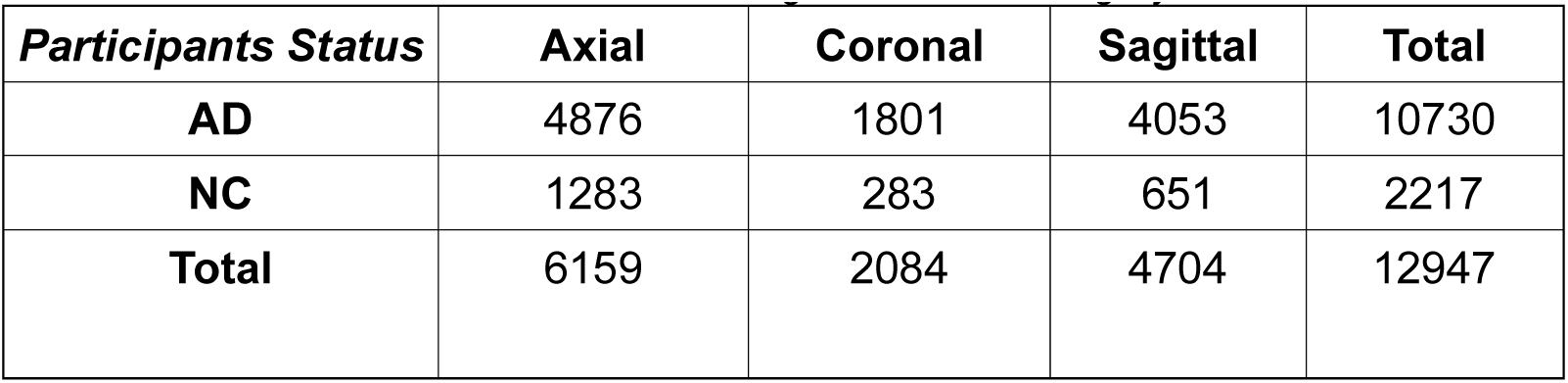
The exact counts of images for each category of AlzaSet.

| **Participants Status** | **Axial** | **Coronal** | **Sagittal** | **Total** |
| --- | --- | --- | --- | --- |
| **AD** | 4876 | 1801 | 4053 | 10730 |
| **NC** | 1283 | 283 | 651 | 2217 |
| **Total** | 6159 | 2084 | 4704 | 12947 |

The dataset was split into training, validation and test subsets to optimize classification algorithm performance. The details are presented in Table 3.

**Table 3.** Distribution of splitted dataset.

| **Axial** | **Train** | **Validation** | **Test** | **Coronal** | **Train** | **Validation** | **Test** | **Sagittal** | **Train** | **Validation** | **Test** |
| --- | --- | --- | --- | --- | --- | --- | --- | --- | --- | --- | --- |
| **AD** | 3459 | 954 | 463 | **AD** | 1247 | 449 | 184 | **AD** | 2823 | 826 | 404 |
| **NC** | 913 | 240 | 130 | **NC** | 184 | 66 | 32 | **NC** | 408 | 160 | 83 |

### 3.2. Data Preprocessing and Augmentation Strategy

To mitigate class imbalance between Alzheimer’s disease (AD) and normal control (NC) images in the AlzaSet dataset, we applied partial data augmentation exclusively to the underrepresented NC cohort. Geometric transformations—including 5% horizontal and vertical translations, 5° rotations, and 0.05× zoom scaling—were employed to synthetically expand the NC subset and approximate numerical parity with AD cases. Representative augmented images from the NC group are illustrated in Figure 1.

**Figure 1.**
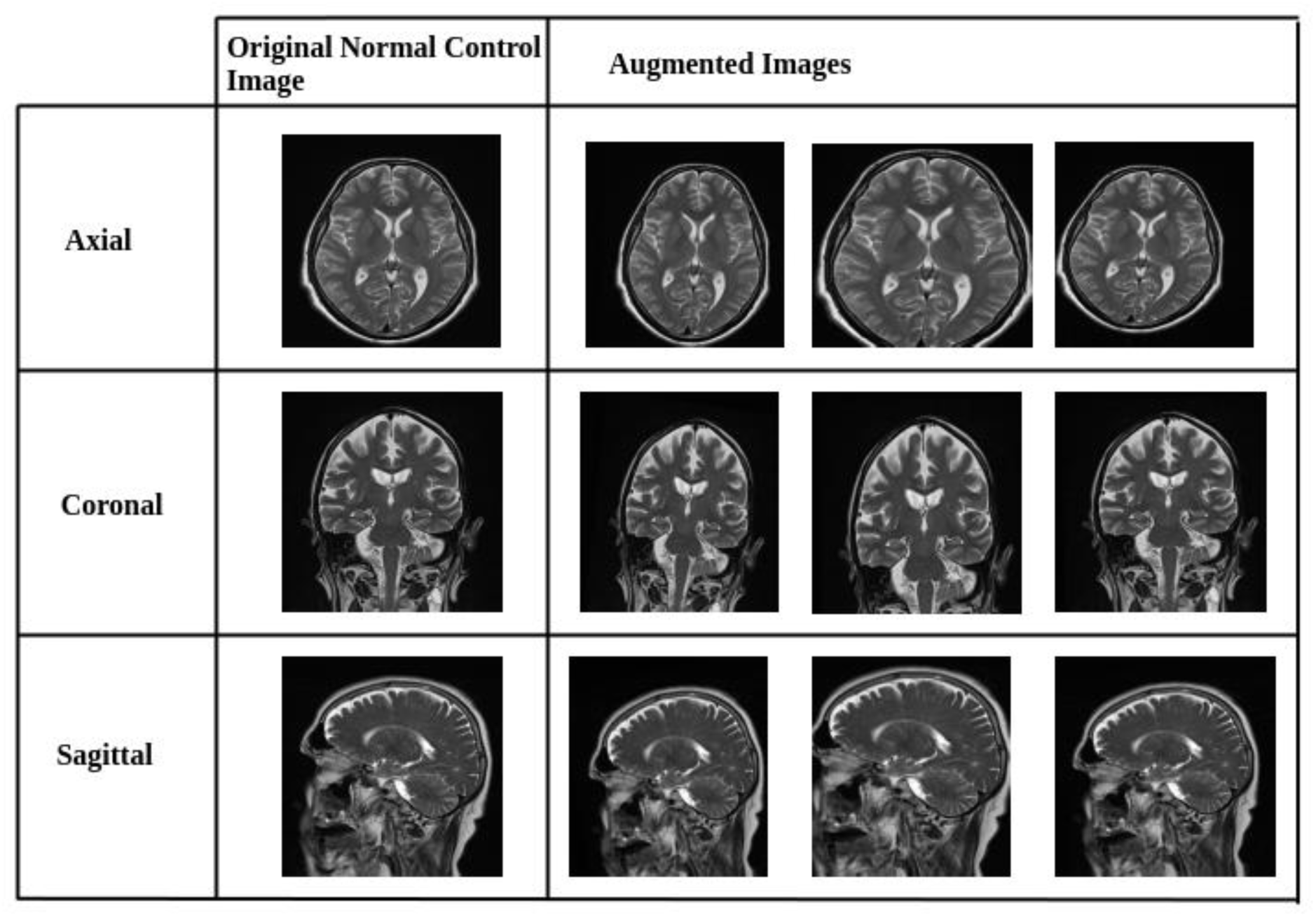
Representative Data Augmentation Pipeline for Normal Control (NC) Class. Sample images from the NC cohort after applying geometric data augmentation techniques. Transformations include horizontal and vertical shifts (±5%), 5-degree rotations, and 0.05× zoom scaling. These augmentations were applied to address class imbalance by synthetically expanding the NC dataset, thereby improving model generalization and reducing overfitting.

Following augmentation, all images were resized to a standardized input dimension of 224 × 224 pixels. Pixel intensities were normalized from the original [0, 255] grayscale range to a floating-point range of [0, 1] to improve numerical stability and reduce computational complexity during model training. The final post-augmentation distribution of the dataset is summarized in Table 4.

**Table 4.**
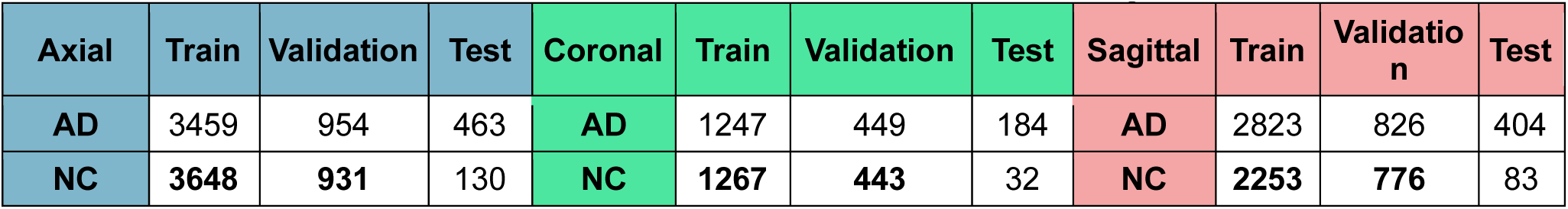
Distribution of AlzaSet after data balancing.

## 4.0 Model Architecture and Implementation

### 4.1. Transfer Learning and Feature Extraction Strategy Using ConvNeXt Architecture

Convolutional neural networks (CNNs) extract higher-level abstract features with hierarchical layers. The initial convolutional layers tend to learn the low-level features such as edges, texture, and gradients that have high domain-transferability even in training on natural image datasets. For medical imaging, transfer learning using feature extraction is a good way of avoiding overfitting as well as enhancing generalizability in the case of limited amounts of data available. With the pretrained convolutional layers frozen, the models retain representations that are generally applicable but fine-tune the classification layers with task-specific data [23].

We utilized the first three blocks of the ConvNeXt model as a pre-trained fixed feature extractor in this work. The pre-trained weights on ImageNet were not changed while the remaining layers were tuned and optimized for AD classification. As observed from Figure 2, the extracted feature maps were input to a customized classification head trained to identify neurodegenerative patterns within structural MRI scans.

**Figure 2.**
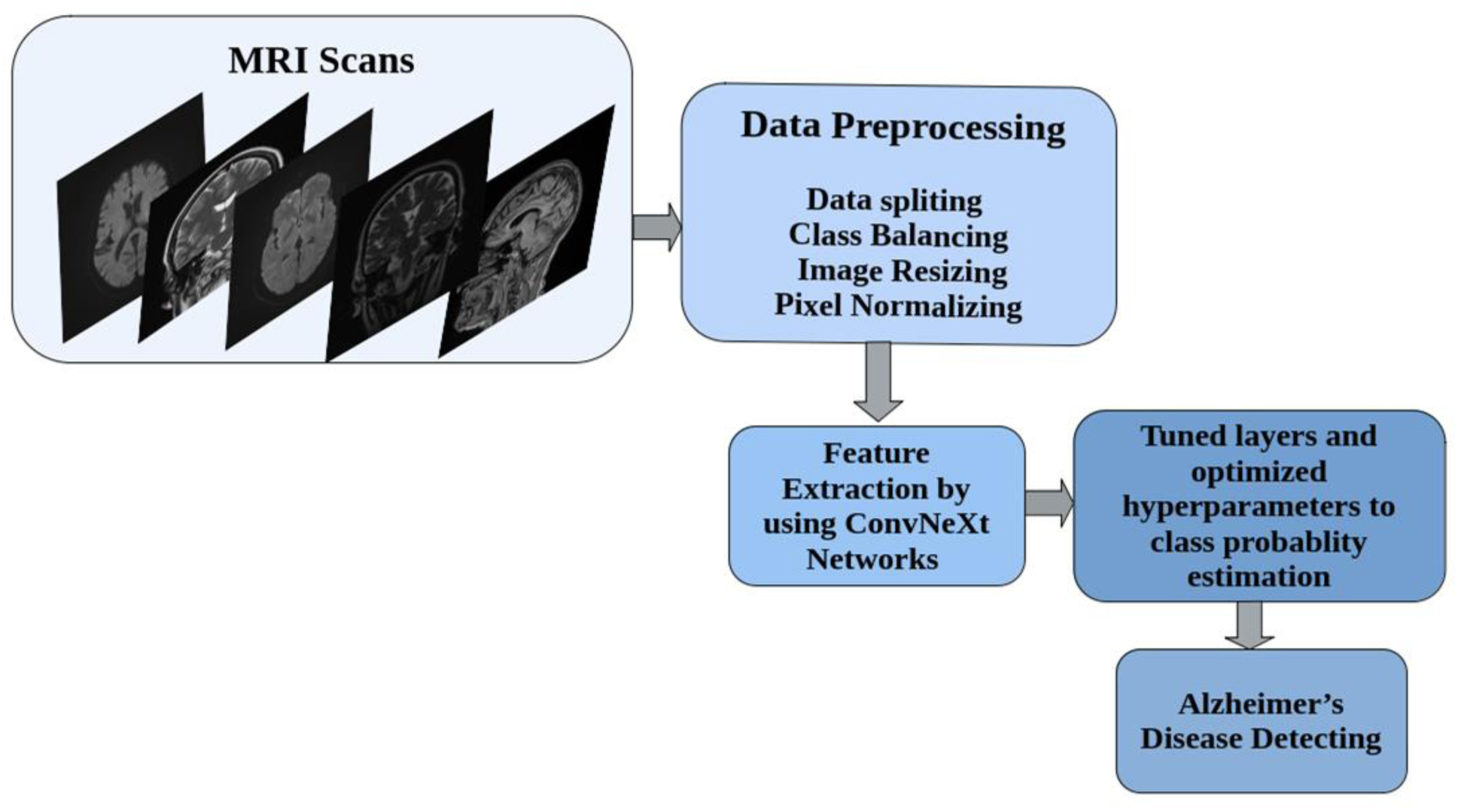
End-to-End Workflow for MRI-Based Alzheimer’s Disease Classification Using ConvNeXt. Overview of the full pipeline for classifying Alzheimer’s disease from structural MRI data. Raw MRI scans in axial, coronal, and sagittal planes undergo a preprocessing stage involving data splitting, class balancing through augmentation, image resizing, and pixel normalization. Preprocessed images are passed through frozen convolutional blocks of ConvNeXt networks for feature extraction. The extracted features are then processed by a set of task-specific layers with optimized hyperparameters to generate final class probability estimates for Alzheimer’s disease diagnosis.

### 4.2. ConvNeXt-Based Model Design Architecture

ConvNeXt is a multi-stage convolutional neural network that integrates design principles from ResNet and Swin Transformer architectures, but is built entirely with convolutional operations. It comprises four consecutive stages and incorporates substantial improvements, larger convolutional kernels (7×7), depthwise separable convolutions, and residual connections, that boost its receptive field while maintaining computational efficiency [13]. ConvNeXt employs non-overlapping convolutional blocks with stride-adjusted kernels, and at its core architectural building block are revamped residual blocks that adopt Transformer-inspired improvements without sacrificing the simplicity of conventional ConvNeXts.

In this study, we developed modified ConvNeXt-Tiny, ConvNeXt-Small, and ConvNeXt-Base architectures, using the original pretrained models as backbones and adding custom adaptations for medical imaging classification:

- **ConvNeXt-Tiny** consists of four stages with [3, 3, 9, 3] blocks, respectively. We preserved the first 125 layers and truncated the final 25 layers to prevent overfitting and reduce model complexity. Each block implements the sequence: **Input** → **LayerNorm** → **PWConv** (1×1) → **GELU** → **DWConv** (7×7) → **PWConv** (1×1) **→ Residual Add**, where **DWConv** denotes depthwise convolution and **PWConv** denotes pointwise convolution.
- **ConvNeXt-Small** shares the same block structure as Tiny but incorporates deeper feature extraction in Stage 3, with [3, 3, 27, 3] blocks. We retained the first 240 layers and removed the final 50 layers for this configuration. Furthermore, the deeper feature extraction results from the iterative application of blocks rather than being an inherent component of the architecture itself.
- **ConvNeXt-Base** expands upon the Small variant by increasing channel widths for greater representational capacity. We extracted features from the first 245 layers, truncating the final 45 layers.

As illustrated in **Figure 3**, each ConvNeXt-based backbone generates a high-dimensional feature map, which is subsequently passed through a custom classification head consisting of the following components:

1. **MaxPooling2D** – A spatial pooling operation (e.g., 2×2) that downsamples the feature map by selecting the maximum activation within each region. This reduces dimensionality, accelerates computation, and mitigates overfitting by emphasizing salient features [24].
2. **Dropout (rate = 0.5)** – A regularization technique that randomly deactivates 50% of neurons during training, improving generalization by preventing co-adaptation of feature detectors [25].
3. **SeparableConv2D** – Implements a two-step process: (i) **depth-wise convolution**, which applies a spatial filter independently to each input channel, followed by (ii) **pointwise convolution** (1×1) to mix inter-channel information. This architecture offers significant computational savings while retaining competitive performance [26].
4. **Batch Normalization** – Normalizes each feature map by applying consistent mean and variance across spatial dimensions. This stabilizes and accelerates training, especially in deep networks [27].
5. **GELU Activation** – The Gaussian Error Linear Unit (GELU) activation, defined as:

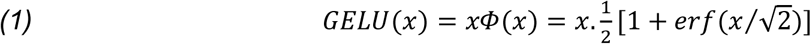

*where*Φ(x)*represents the standard Gaussian cumulative distribution function and erf denotes the error function. The GELU outperforms other activation functions such as ReLU and ELU across multiple tasks, as demonstrated in the experiments* [28].
6. **GlobalAveragePooling2D** – Reduces each feature map to a single scalar by averaging across all spatial dimensions (height and width), effectively summarizing global context while minimizing parameter count [29].
7. **Dense Layer with Softmax Activation** – A fully connected output layer containing two neurons, representing the binary classes (AD and NC). The softmax activation function generates normalized probability scores for final classification.

**Figure 3.**
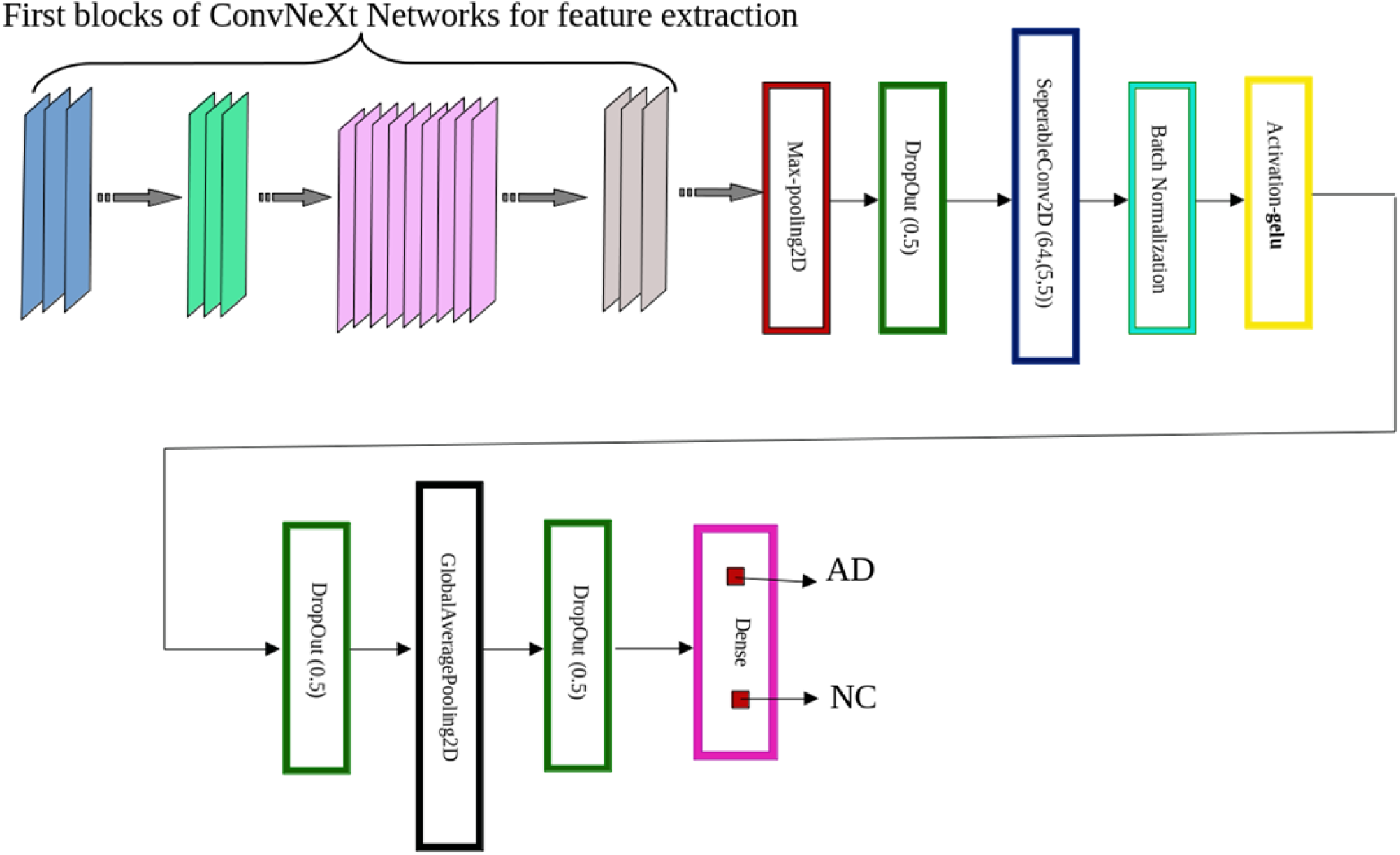
Architecture of the Proposed ConvNeXt-Based Alzheimer’s Disease Classifier. Schematic representation of the model architecture used for Alzheimer’s disease detection. The initial layers of a pretrained ConvNeXt variant (Tiny, Small, or Base) are used for feature extraction. These are followed by a custom classification head comprising a MaxPooling2D layer, Dropout (rate = 0.5), SeparableConv2D, Batch Normalization, and GELU activation. The output is further processed by GlobalAveragePooling2D, additional Dropout layers, and a final Dense layer with two output neurons activated by softmax to classify inputs as either Alzheimer’s disease (AD) or normal control (NC).

### 4.3. Training Configuration and Hyperparameters

To optimize model performance, we carefully selected hyperparameters based on standard practices in deep learning and task-specific requirements. Table 5. summarizes the key training hyperparameters and their configurations.

**Table 5.** Model Training Hyperparameters.

| Hyperparameter | Value | Description |
| --- | --- | --- |
| Batch Size | 32 | Chosen due to GPU memory constraints. |
| Loss Function | Categorical Cross-Entropy | Selected due to its ability to measure the dissimilarity between predicted probabilities and true class labels, thereby simplifying optimization and enhancing model performance. |
| Epochs | 35 | Selected through validation(tested epochs: 20, 35,50,100) with 35 demonstrating model convergence. |
| Optimizer | Adam | Based on its demonstrated advantages in both optimization efficiency and generalization performance. |
| Learning Rate | 1e-4 | The tested range (1e-3 to 1e-5) and final selection 1e-4. |

## 5. Performance Evaluation

### 5.1. Classification Metrics

To validate the correctness of a classification task, we compute four measures:

1. ***True Positives (tp)*:** Correctly identified instances of the class.
2. ***True Negatives (tn):*** Correctly rejected instances that don’t belong to the class.
3. ***False Positives (fp)*:** Instances wrongly classified as part of the class.
4. ***False Negatives (fn):*** Actual class instances that were missed. These measurements contribute **Confusion Matrix** that presented in Table 6. [30]

**Table 6.** Confusion Matrix for Classification.

|  | Predicted Positives | Predicted Negatives |
| --- | --- | --- |
| Actual Positives | tp | fn |
| Actual Negatives | fp | tn |

We evaluate our proposed model performance using standard metrics: Accuracy, Precision, Recall (Sensitivity), F1-Score and AUC, that computed based on confusion matrix.

**Accuracy,** computed as the proportion of correctly predicted instances relative to the total predictions, represents a fundamental metrics for evaluating model efficiency in classification tasks. Accuracy is obtained from the Equation 2. [31]

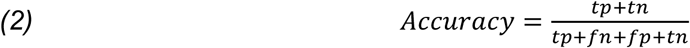

**Precision** calculated using Equation 3. represents the proportion of correctly predicted positive instances among all positive predictions. (i.e. cases identified as AD in our study).[32]

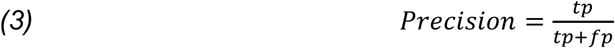

**Recall (Sensitivity),** computed as proportion of correctly AD predicted among all actual AD cases.(Equation 4.)

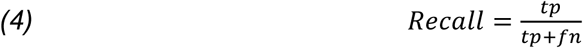

**F1-Score,** (Equation 5.) provides the harmonic mean of Precision and Recall.

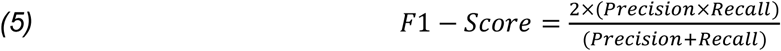

The **Area Under the ROC Curve (AUC)** is a scalar metric that measures the model’s ability to discriminate between positive (AD) and negative (NC) classes. [33]

## 6.0 Results

### 6.1. Model Performance Across Anatomical Planes

This section presents a comprehensive analysis of our proposed architecture’s performance across multiple ConvNeXt configurations(Tiny, Small, and Base variants) using the aforementioned evaluation metrics. The benchmark results, obtained on our newly introduced AlzaSet Dataset. Classification results across anatomical planes are presented as follows:

Table 7. (**Axial** Plane): Diagnostic accuracy and evaluation metrics for transverse view analysis.

**Table 7.** Test Set Performance Metrics for Axial Plane Images.

| **Axial** | **Accuracy** | **Loss** | **Precision Macro Avg** | **Recall Macro Avg** | **F1-Score Macro Avg** | **AUC** |
| --- | --- | --- | --- | --- | --- | --- |
| **ConvNeXt-Tiny** | 0.9303 | 0.2590 | 0.9400 | 0.8500 | 0.8800 | 0.8470 |
| **ConvNeXt-Small** | 0.8877 | 0.3388 | 0.9200 | 0.7500 | 0.8000 | 0.7480 |
| **ConvNeXt-Base** | 0.9208 | 0.2176 | 0.9500 | 0.8300 | 0.8700 | 0.8290 |

Table 8. (**Coronal** Plane): Diagnostic accuracy and evaluation metrics for frontal view analysis.

**Table 8.** Test Set Performance Metrics for Coronal Plane Images.

| Coronal | Accuracy | Loss | Precision Macro Avg | Recall Macro Avg | F1-Score Macro Avg | AUC |
| --- | --- | --- | --- | --- | --- | --- |
| ConvNeXt-Tiny | 0.9896 | 0.1015 | 0.9900 | 0.9700 | 0.9800 | 0.9680 |
| ConvNeXt-Small | 0.9579 | 0.1339 | 0.8900 | 0.9600 | 0.9200 | 0.9620 |
| ConvNeXt-Base | 0.9837 | 0.0924 | 0.9600 | 0.9900 | 0.9700 | 0.9920 |

Table 9. (**Sagittal** Plane): Diagnostic accuracy and evaluation metrics for lateral view analysis.

**Table 9.** Test Set Performance Metrics for Sagittal Plane Images.

| Sagittal | Accuracy | Loss | Precision Macro Avg | Recall Macro Avg | F1-Score Macro Avg | AUC |
| --- | --- | --- | --- | --- | --- | --- |
| ConvNeXt-Tiny | 0.9077 | 0.3011 | 0.9200 | 0.7400 | 0.8000 | 0.7430 |
| ConvNeXt-Small | 0.8754 | 0.3208 | 0.8000 | 0.6800 | 0.7200 | 0.6830 |
| ConvNeXt-Base | 0.9209 | 0.1791 | 0.9300 | 0.7800 | 0.8300 | 0.7790 |

**Table 10.** Performance of Pre-trained Networks on Axial Plane Images of AlzaSet.

| **Axial** | **Accuracy** | **Loss** | **Precision**<br>**Macro Avg** | **Recall**<br>**Macro Avg** | **F1-Score**<br>**Macro Avg** | **AUC** |
| --- | --- | --- | --- | --- | --- | --- |
| **VGG16** | 0.8799 | 0.2771 | 0.9000 | 0.7400 | 0.7800 | 0.7370 |
| **VGG19** | 0.8734 | 0.3564 | 0.9100 | 0.7100 | 0.7600 | 0.7090 |
| **InceptionV3** | 0.8954 | 0.2006 | 0.9000 | 0.7900 | 0.8300 | 0.7910 |
| **DenseNet121** | 0.9043 | 0.2606 | 0.8900 | 0.8000 | 0.8300 | 0.8002 |

**Table 11.** Performance of Pre-trained Networks on Coronal Plane Images of AlzaSet.

| **Coronal** | **Accuracy** | **Loss** | **Precision**<br>**Macro Avg** | **Recall**<br>**Macro Avg** | **F1-Score**<br>**Macro Avg** | **AUC** |
| --- | --- | --- | --- | --- | --- | --- |
| **VGG16** | 0.9722 | 0.1780 | 0.9800 | 0.9400 | 0.9000 | 0.9030 |
| **VGG19** | 0.9678 | 0.2133 | 0.9400 | 0.9300 | 0.9300 | 0.9270 |
| **InceptionV3** | 0.9513 | 0.2966 | 0.8900 | 0.9500 | 0.9100 | 0.9460 |
| **DenseNet121** | 0.9415 | 0.2910 | 0.8700 | 0.9400 | 0.9000 | 0.9410 |

**Table 12.** Performance of Pre-trained Networks on Sagittal Plane Images of AlzaSet Performance of Pre-trained Networks on Sagittal Plane Images of AlzaSet.

| **Sagittal** | **Accuracy** | **Loss** | **Precision**<br>**Macro Avg** | **Recall**<br>**Macro Avg** | **F1-Score**<br>**Macro Avg** | **AUC** |
| --- | --- | --- | --- | --- | --- | --- |
| **VGG16** | 0.8963 | 0.2573 | 0.9400 | 0.6900 | 0.7500 | 0.6930 |
| **VGG19** | 0.8731 | 0.2616 | 0.9300 | 0.6300 | 0.6700 | 0.6330 |
| **InceptionV3** | 0.9204 | 0.2003 | 0.9100 | 0.7900 | 0.8300 | 0.7880 |
| **DenseNet121** | 0.8936 | 0.2874 | 0.8200 | 0.7800 | 0.7900 | 0.7760 |

We compared the performance of three versions of ConvNeXt—Tiny, Small, and Base—on the AlzaSet dataset in terms of classification performance across axial, coronal, and sagittal anatomical views. Overall, with all views taken together, the ConvNeXt-Base model performed better than its smaller counterparts uniformly and had the best values of accuracy, precision, recall, F1-score, and AUC. For the axial plane, ConvNeXt-Base scored 92.08% accuracy and an AUC of 0.829, lagging behind ConvNeXt-Tiny slightly in accuracy but showing better balance between precision and recall. ConvNeXt-Small demonstrated the weakest performance among the three variants, likely due to its limited depth and a suboptimal balance between model complexity and overfitting risk.

Sagittal plane performance also maintained a similar trend, where ConvNeXt-Base achieved a 92.09% classification accuracy and an AUC of 0.779. Tiny trailed closely behind, followed by the Small variant with lower classification accuracy. ConvNeXt-Base delivered its highest performance in the coronal plane, achieving a 98.37% classification accuracy and 0.992 AUC. ConvNeXt-Tiny also delivered an excellent performance in this view, achieving a close to 99% accuracy. These findings are a validation set for diagnostic utility of coronal images, which provide superior anatomical definition of medial temporal lobe structures engaged early in AD pathology. They include the hippocampus and the parahippocampal gyrus.

### 6.2. Comparative Benchmarking Against Established Models

To place our results in context, we compared ConvNeXt-Base with four well-known CNN architectures—VGG16, VGG19, InceptionV3, and DenseNet121—each with the same preprocessing and hyperparameter training. ConvNeXt-Base outperformed these established models in all the anatomical planes. In axial view, ConvNeXt-Base performed better than DenseNet121 by a good margin in both AUC (0.829 vs. 0.8002) and F1-score (0.870 vs. 0.830). In coronal view, ConvNeXt-Base recorded the highest AUC (0.992) and F1-score (0.970) among all models tested. In sagittal plane, ConvNeXt-Base also performed best in the majority of the metrics, demonstrating how it is robust across orientations. The superiority of ConvNeXt models is also better highlighted in Figures 5 to 7, comparing performance on key classification metrics, and demonstrating clearly their discriminability and generalizability.

Confusion matrix plots and ROC curves (Figure 4) also strongly support the stability of the ConvNeXt-Base model. On the coronal test set, it had low misclassification and virtually perfect class separation, again strongly supporting the conclusion that architectural depth as well as anatomical orientation both have key roles to play in enhancing diagnostic accuracy.

**Figure 4.**
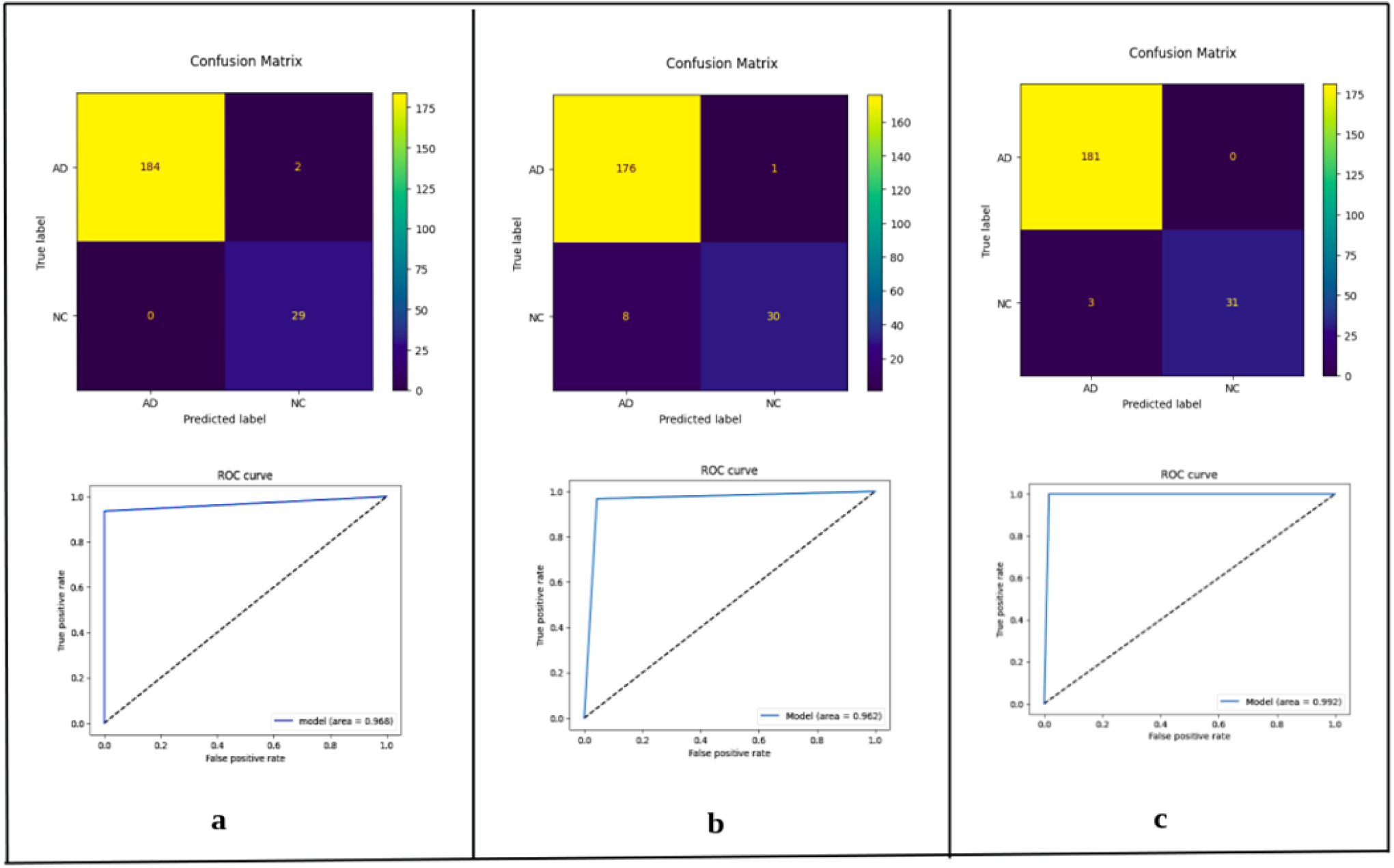
Confusion Matrices and ROC Curves for ConvNeXt Variants on Coronal Plane Test Images. Visual evaluation of classification performance on coronal MRI slices using the three ConvNeXt variants: (a) ConvNeXt-Tiny, (b) ConvNeXt-Small, and (c) ConvNeXt-Base. The top row depicts confusion matrices, where yellow cells indicate correct classifications and darker cells indicate misclassifications. The bottom row presents the corresponding Receiver Operating Characteristic (ROC) curves, with Area Under the Curve (AUC) values of 0.968, 0.962, and 0.992 for Tiny, Small, and Base respectively. The results illustrate the high discriminative power of ConvNeXt models on coronal images, particularly the Base variant, which demonstrates near-perfect separation between Alzheimer’s disease (AD) and normal control (NC) classes.

**Figure 5.**
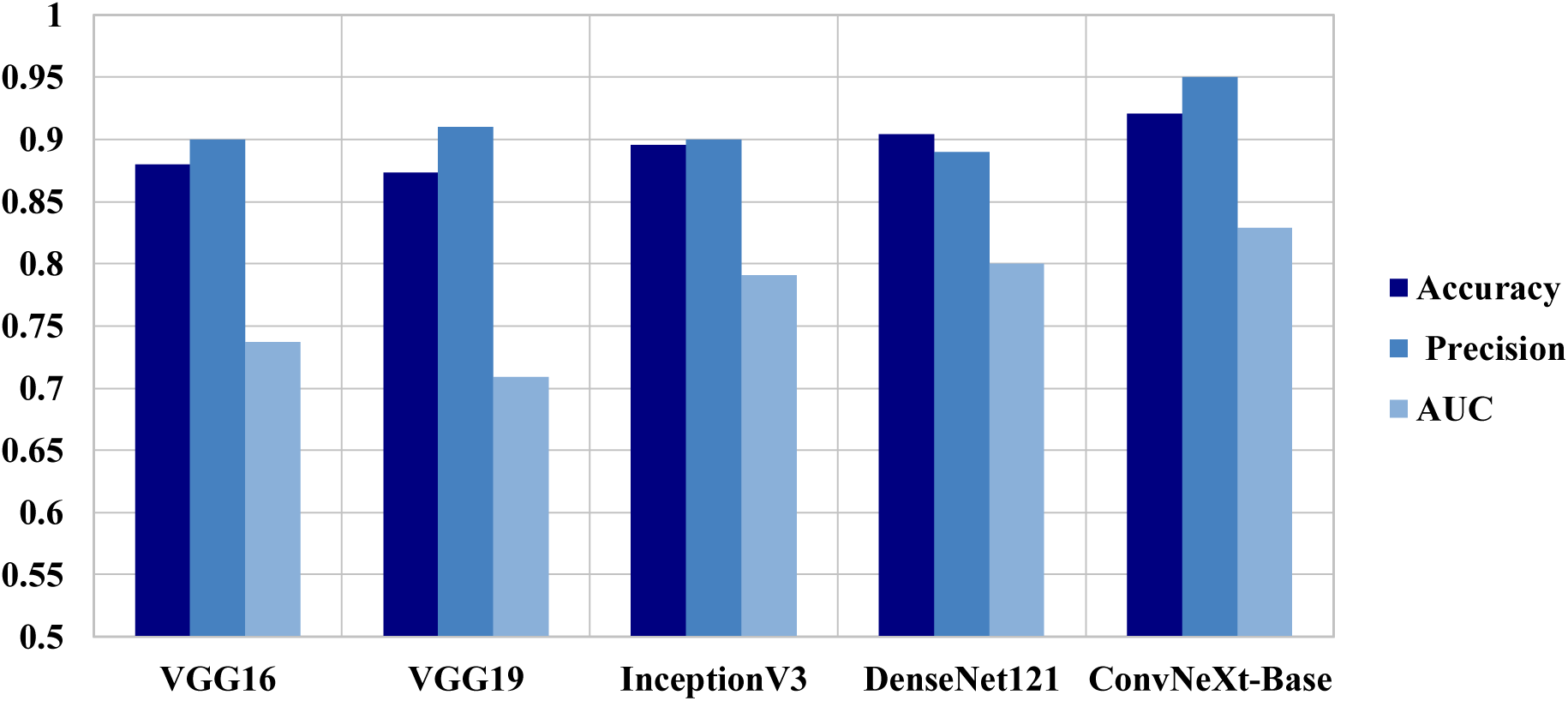
Comparative Performance of ConvNeXt Versus Pre-trained Models Across Axial Plane Images. Bar plot comparing the classification performance of ConvNeXt-Base against standard pre-trained convolutional models (VGG16, VGG19, InceptionV3, and DenseNet121) on axial MRI slices from the AlzaSet dataset. Metrics displayed include Accuracy, Precision, and Area Under the Curve (AUC). ConvNeXt-Base outperforms all benchmark models across all three evaluation metrics, highlighting its superior capacity for learning discriminative features relevant to Alzheimer’s pathology in axial views.

**Figure 6.**
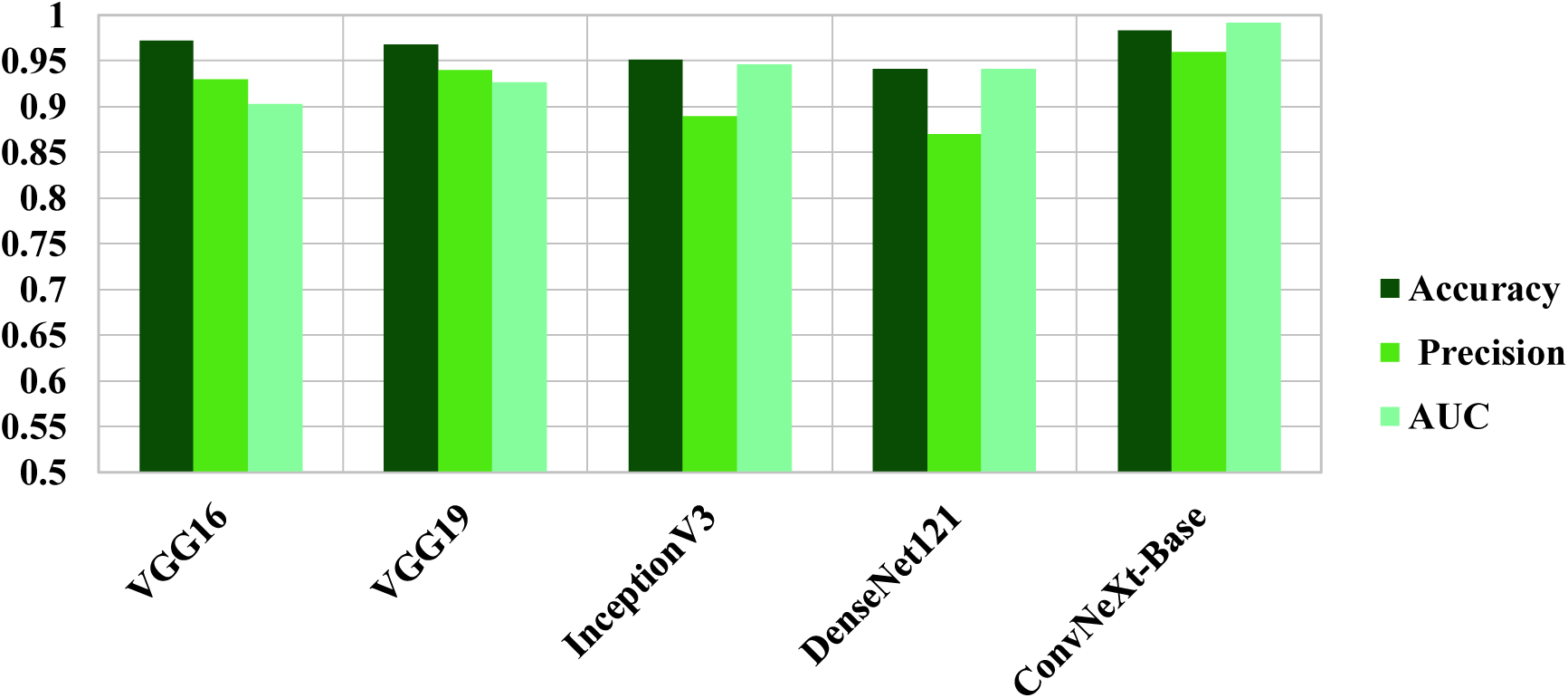
Comparative Performance of ConvNeXt versus Pre-trained Models Across Coronal Plane Images. Bar chart illustrating the performance comparison of ConvNeXt-Base against four widely adopted pre-trained CNN models—VGG16, VGG19, InceptionV3, and DenseNet121—on coronal MRI slices from the AlzaSet dataset. Evaluation metrics include Accuracy (dark green), Precision (medium green), and Area Under the ROC Curve (AUC; light green). ConvNeXt-Base demonstrates superior performance across all three metrics, particularly in AUC, underscoring the enhanced discriminatory power of coronal slices and the architectural advantage of ConvNeXt for capturing Alzheimer-related features in medial temporal lobe structures.

**Figure 7.**
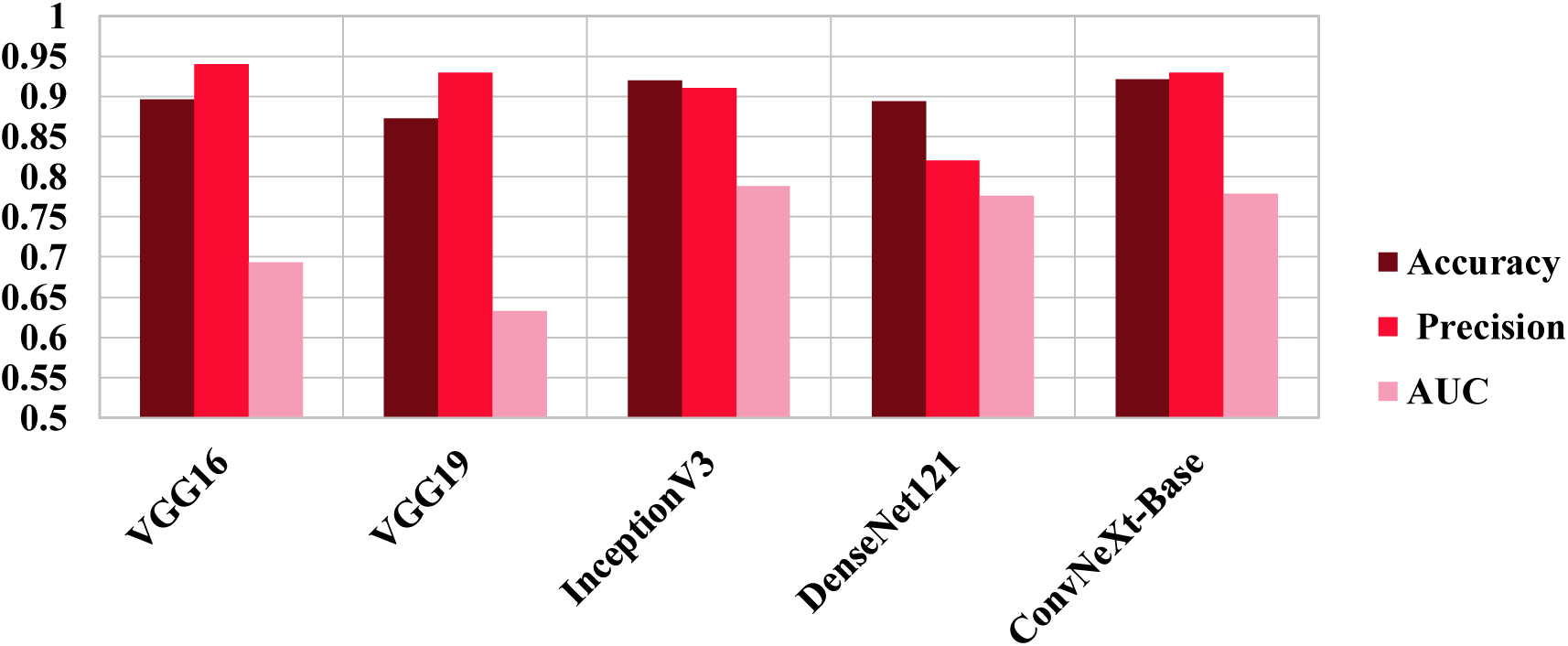
Comparative Performance of ConvNeXt-Base Versus Pretrained CNN Models on Sagittal MRI Plane Images. Performance comparison of ConvNeXt-Base against VGG16, VGG19, InceptionV3, and DenseNet121 on sagittal MRI slices from the AlzaSet dataset. ConvNeXt-Base achieved the highest accuracy (92.1%) and precision (94%), with strong AUC performance (0.779), highlighting its superior ability to detect Alzheimer’s disease in lateral brain views despite anatomical limitations.

## 7.0 Discussion

### 7.1. Interpretation of Findings and Architectural Insights

Our results solidly establish the effectiveness of ConvNeXt models for AD prediction from clinical images, particularly after training on high-quality datasets such as AlzaSet. ConvNeXt not only enjoys better classification performance relative to traditional CNNs but also offers deployment advantages in practice with its modularity, efficiency, and interoperability with modern training schemes.

The better performance of ConvNeXt-Base than both older CNNs and the smaller ConvNeXt models reveals that deeper representations of features, appropriately regularized, are required for the representation of the subtle structural changes characteristic of Alzheimer’s disease. Furthermore, our plane-wise analysis revealed that coronal slices consistently yielded the maximum diagnostic accuracy in all models, supporting previous radiological evidence emphasizing medial temporal lobe visualization as key.

### 7.2. Integration with Prior Literature

These findings are also supported when compared with more recent work by Techa et al., where they built an ensemble structure for AD classification using ConvNeXt, on a Kaggle dataset [34]. While their model achieved high accuracy (92.2%) using stacking classifiers, their pipeline introduced an extra layer of complexity by combining multiple machine learning classifiers (i.e., SVM, MLP, Decision Trees) post-feature extraction. Conversely, our effort utilized an end-to-end deep learning system, streamlining the pipeline and reducing inference overhead. Our model’s emphasis on orientation-specific evaluation (axial, coronal, sagittal) and use of an expert-annotated clinical dataset further distinguishes our approach both methodologically and in terms of clinical relevance. In addition, a recent study by Moscoso et al. (2025) in JAMA highlights the prognostic value of tau PET imaging, demonstrating that individuals who are both Aβ- and tau-positive have a significantly elevated risk of clinical conversion from preclinical to MCI or dementia [35]. This favors the usefulness of multimodal approaches, such as combining deep learning on structural MRI with PET-based biomarkers, to stage AD pathology earlier and more precisely. Such biomarker stratification incorporated into model training pipelines can enhance prognostic significance and clinical usefulness.

Another unique feature of the current work is in the target classification task. While Techa et al. addressed multi-class classification across different AD stages (non-demented, very mild, mild, moderate), our methodology aimed at binary classification (AD vs. NC), which is more clinically pertinent in the immediate sense for early screening and detection. In addition, AlzaSet’s dataset, developed in cooperation with clinical site’s neuroradiologists, provides higher consistency and diagnostic validity over the publicly available, unevenly annotated Kaggle dataset. These differences highlight the translational potential of our model.

### 7.3. Clinical Implications and Future Directions

The clinical applicability of our results is significant. Notably, the ConvNeXt-Base model demonstrates classification accuracy on par with that of expert radiologists for binary diagnosis tasks, particularly for the coronal plane. In addition, model variants, Tiny and Base, offer computational cost vs. accuracy trade-offs for flexible deployment over a broad range of clinical settings. Compared to transformer-based or ensemble-based methods, ConvNeXt’s purely convolutional architecture maintains spatial locality, enabling easier application with post hoc interpretability techniques like Grad-CAM.

Furthermore, while we did not yet apply formal interpretability techniques (e.g., Grad-CAM) to spatially localize activation maps, the model’s superior performance on coronal slices strongly implicates disease-relevant changes in the medial temporal lobe. Specifically, ConvNeXt-Base could have learned to recognize early patterns of atrophy in the hippocampus, entorhinal cortex, and parahippocampal gyrus, regions frequently involved in prodromal and mild Alzheimer’s disease. These regions exhibit volume loss, thinning of the cortex, and architectural distortion on T1-weighted MRI and are optimally viewed in coronal planes. The use of SeparableConv2D and GlobalAveragePooling also enabled the model to compact localized structural abnormalities into informative features, which optimized its capability for classifying early disease stages. Subsequent studies using interpretability tools will allow a more precise ascription of these spatial features, possibly leading to earlier diagnostic triaging in the clinic.

Yet, this study is not without limitations. Our database, although well-annotated, is from a single geographic site and hence cannot be expected to generalize to populations with varied demographics, or diverse imaging protocols. Moreover, we employed only T1-weighted MRI data; multimodal imaging inputs such as PET, diffusion-weighted MRI, or resting-state fMRI might serve to enhance model discriminability. Lastly, the binary nature of our classification task limits applicability to staging or longitudinal prediction of disease progression.

One strong limitation of our study is the timing of image acquisition. All MRIs utilized in model training were obtained after clinical diagnosis of Alzheimer’s disease, and thus, imaging reflects relatively advanced disease stages already detected by treating physicians. Even though our model performs well in differentiating AD from normal controls in this instance, the maximum clinical utility would be to detect AD prior to clinical diagnosis, particularly during the stage of preclinical or prodromal disease when intervention would be more useful. Future studies should attempt to train and validate models on imaging acquired before symptom onset or on early cognitive change, preferably supplemented with neuropsychological test scores, fluid biomarkers, or PET images to add specificity. Moreover, higher resolution characterization of the imaging features driving classification, through activation mapping or voxel-wise morphometry, will be required in order to unequivocally characterize the earliest radiographic markers of disease that may guide preemptive diagnosis and treatment.

Future directions involve increasing the size of our dataset by including images from other institutions and multi-modal neuroimaging data. Addition of attention-based architectures or hybrid CNN-transformer architectures could also provide increases in performance. Moreover, exploration of using self-supervised pretraining over large, unlabeled neuroimaging datasets could reduce data sparsity and further enhance generalizability.

### 7.4. Conclusion

Overall, this paper presents an efficient, scalable approach to automating Alzheimer’s disease diagnosis on 2D MRI images with ConvNeXt-based deep learning. The presentation of the AlzaSet dataset, exploration of anatomical orientation, and comparative benchmarking against baseline CNNs all contribute to highlighting ConvNeXt-Base architecture as a potentially optimal solution for clinical decision support in AD diagnostics. In contrast to ensemble-based pipelines, our minimalistic architecture maintains accuracy while facilitating model interpretability and efficiency. Following further validation and scale-up, this platform can assist in real-time clinical deployment, especially in resource-constrained environments, enabling earlier and more equitable diagnosis of Alzheimer’s disease worldwide.

## References

[1] WHO. https://www.who.int/news-room/fact-sheets/detail/dementia. (2025)

[2] Cataldi R, Chowdhary N, Seeher K, Moorthy V, Dua T. A blueprint for the worldwide research response to dementia. Lancet Neurol. 2022;21(8):690–691. doi:10.1016/S1474-4422(22)00269-1

[3] David E. Bloom, David Canning, Alyssa Lubet. (2015). “ Global Population Aging: Facts, Challenges, Solutions & Perspectives”. Dædalus, the Journal of the American Academy of Arts & Sciences. DOI: 10.1162/DAED_a_00332

[4] Braak H, Braak E, Bohl J. Staging of Alzheimer-related cortical destruction. Eur Neurol. 1993;33(6):403–408. doi:10.1159/000116984

[5] Clifford R. Jack Jr, Heather J. Wiste, Prashanthi Vemuri. (2010).“Brain beta-amyloid measures and magnetic resonance imaging atrophy both predict time-to-progression from mild cognitive impairment to Alzheimer’s disease”. Brain: A Journal of Neurology. DOI : 10.1093/brain/awq277

[6] Ajam Oughli H, Siddarth P, Lavretsky H, et al. Peripheral Alzheimer’s disease biomarkers are related to change in subjective memory in older women with cardiovascular risk factors in a trial of yoga vs memory training. Can J Psychiatry. 2025. doi:10.1177/07067437251343291

[7] Abikenari M, Jain B, Xu R, et al. Bridging imaging and molecular biomarkers in trigeminal neuralgia: Toward precision diagnosis and prognostication in neuropathic pain. Med Res Arch. 2025;13(5). doi:10.18103/mra.v13i5.6605

[8] Siddarth P, Abikenari M, Grzenda A, et al. Inflammatory markers of geriatric depression response to Tai Chi or health education adjunct interventions. Am J Geriatr Psychiatry. 2023;31(1):22–32. doi:10.1016/j.jagp.2022.08.004

[9] Geert Litjens, Thijs Kooi, Babak Ehteshami Bejnordi. (2017). “ A Survey on Deep Learning in Medical Image Analysis”. Medical Image Analysis. DOI: 10.1016/j.media.2017.07.005

[10] Jason Yosinski, Jeff Clune, Yoshua Bengio. (2014). “How transferable are features in deep neural networks?”. Proceedings of the 28th International Conference on Neural Information Processing Systems - Volume 2. DOI: 10.48550/arXiv.1411.1792

[11] Matthew D. Zeiler, Rob Fergus. (2014). “Visualizing and Understanding Convolutional Networks”. European Conference on Computer Vision. DOI: 10.48550/arXiv.1311.2901

[12] Hoo-Chang Shin, Holger R. Roth, Mingchen Gao. (2016). “Deep Convolutional Neural Networks for Computer-Aided Detection: CNN Architectures, Dataset Characteristics and Transfer Learning”. IEEE Transactions on Medical Imaging. DOI: 10.1109/TMI.2016.5291612

[13] Zhuang Liu, Hanzi Mao, Chao-Yuan Wu. (2022).“A ConvNet for the 2020s”. Proc. IEEE/CVF Conf. Comput. DOI: 10.48550/arXiv.2201.03545

[14] Siyuan Lu, Zhihai Lu, Yu-Dong Zhang. (2018). “Pathological Brain Detection based on AlexNet and Transfer Learning”. Journal of Computational Science. DOI: 10.1016/j.jocs.2018.11.008

[15] Naimul Mefraz Khan, Nabila Abraham, Marcia Hon. (2019). “Transfer Learning With Intelligent Training Data Selection for Prediction of Alzheimer’s Disease”. IEEE Access. DOI : 10.1109/ACCESS.2019.2920448

[16] Loris Nanni, Matteo Interlenghi, Shery Brahman. (2020). “Comparison of Transfer Learning and Conventional Machine Learning Applied to Structural Brain MRI for the Early Diagnosis and Prognosis of Alzheimer’s Disease”. Frontiers in Neurology. Dementia and Neurodegenerative Diseases. DOI: 10.3389/fneur.2020.576194

[17] Atif Mehmood, Shuyuan Yang, Zhixi Feng. (2021). “A Transfer Learning Approach for Early Diagnosis of Alzheimer’s Disease on MRI Images. Neuroscience, Elsevier. DOI: 10.1016/j.neuroscience.2021.01.002

[18] Rizwan Khan, Saeed Akbar, Atif Mehmood. (2022). “A transfer learning approach for multiclass classification of Alzheimer’s disease using MRI images”. Frontiers in Neuroscience. DOI: 10.3389/fnins.2022.1050777

[19] Tanjim Mahmud, Koushick Barua, Anik Barua. (2023). “Exploring Deep Transfer Learning Ensemble for Improved Diagnosis and Classification of Alzheimer’s Disease”. Lecture Notes in Computer Science. DOI: 10.1007/978-3-031-43075-6_10

[20] Kishor Kumar Reddy C, Aarti Rangarajan, Deepti Rangarajan. (2024). “A Transfer Learning Approach: Early Prediction of Alzheimer’s Disease on US Healthy Aging Dataset”. Mathematics (ISSN 2227-7390). DOI: 10.3390/math12142204

[21] Zhongyi Hu, Yuhang Wang, Lei Xiao. (2025). “Alzheimer’s disease diagnosis by 3D-SEConvNeXt”. Journal of Big Data. DOI: 10.1186/s40537-025-01088-8

[22] Weikang Jin, Yue Yin, Jing Bai. (2024). “CA-ConvNeXt:Coordinate Attention on ConvNeXt for Early Alzheimer’s disease classification”. Intelligence Science IV. ICIS 2022. IFIP Advances in Information and Communication Technology, vol 659. Springer, Cham. DOI: 10.1007/978-3-031-14903-0_48

[23] Muhamad Angriawan. (2023). “Transfer Learning Strategies for Fine-Tuning Pretrained Convolutional Neural Networks in Medical Imaging”. Research Journal of Computer Systems and Engineering (RJCSE). DOI:10.52710/rjcse.79

[24] Shallu Sharma, Rajesh Mehra. (2019). “Implications of Pooling Strategies in Convolutional Neural Networks: A Deep Insight”. Foundations of Computing and Decision Sciences. Vol. 44 (2019) No. 3. DOI: 10.2478/fcds-2019-0016

[25] G. E. Hinton, N. Srivastava, A. Krizhevsky. (2012). “Improving neural networks by preventing co-adaptation of feature detectors”. DOI: 10.48550/arXiv.1207.0580

[26] Francois Chollet. (2017). “Xception: Deep Learning with Depthwise Separable Convolutions”. IEEE Conference on Computer Vision and Pattern Recognition (CVPR). DOI: 10.1109/CVPR.2017.195

[27] Sergey Ioffe, Christian Szegedy. (2015). “Batch Normalization: Accelerating Deep Network Training by Reducing Internal Covariate Shift”. ICML’15: Proceedings of the 32nd International Conference on International Conference on Machine Learning - Volume 37 Pages 448 – 456. DOI: 10.48550/arXiv.1502.03167

[28] Dan Hendrycks, Kevin Gimpel. (2016). “Gaussian Error Linear Units (GELUs)”. DOI: 10.48550/arXiv.1606.08415

[29] Min Lin, Qiang Chen, Shuicheng Yan. (2013). “Network In Network”. DOI: 10.48550/arXiv.1312.4400

[30] Diederik Kingma, Jimmy Ba. (2014). “Adam: A Method for Stochastic Optimization”. DOI: 10.48550/arXiv.1412.6980

[31] Marina Sokolova, Guy Lapalme. (2009). “A systematic analysis of performance measures for classification tasks”. Information Processing & Management. DOI: 10.1016/j.ipm.2009.03.002

[32] Kohavi, R., Provost, F. (1998). “Glossary of terms. Stanford University Technical Report”. https://ai.stanford.edu/~ronnyk/glossary.html

[33] Tom Fawcett. (2006). “An introduction to ROC analysis”. Pattern Recognition Letters (Volume 27, Issue 8, June 2006). DOI: 10.1016/j.patrec.2005.10.010

[34] Techa, Chaima, et al. “Automated Alzheimer’s disease classification from brain MRI scans using ConvNeXt and ensemble of machine learning classifiers.” International Conference on Soft Computing and Pattern Recognition. Cham: Springer Nature Switzerland, 2022.

[35] Moscoso A, Heeman F, Raghavan S, et al. Frequency and Clinical Outcomes Associated With Tau Positron Emission Tomography Positivity. JAMA. Published online June 16, 2025. doi:10.1001/jama.2025.7817

