## Supplemental Table 2 for "ConvNeXt-Driven Detection of Alzheimer’s Disease: A Benchmark Study on Expert-Annotated AlzaSet MRI Dataset Across Anatomical Planes"

### Supplementary Materials for “ConvNeXt for Alzheimer’s Detection: Benchmarking Modern Architecture on the AlzaSet MRI Dataset”.

This document supplements the main manuscript with:

- Additional Results: Complete per-plane performance metrics, confusion matrices, and ROC curves. (`Supplementary-Materials/TestSet-results’)
- Hardware specifications: All experiments were conducted on Google Colab free and Google Colab pro (Jun-July2024, Order number : COL.3347-8355-8490-08315) with the following resourses. (Table S1)

Table S1: Hardware specifications

| **Component** | **Specification** |
|---|---|
| CPU | Intel Xeon (2.2+ GHz, 2 cores) |
| GPU | Tesla T4 (16GB) \|V100 (16GB) |
| RAM | 12-52GB |

- Table S2: Python Libraries

| **Library** | **Version** |
|---|---|
| TensorFlow | 2.12.1 \| 2.16.1 |
| Keras | 3.3.3 |
| NumPy | 1.26.4 |
| Matplotlib | 3.8.4 |
| Seaborn | 0.13.2 |

- AlzaSet Dataset: The AlzaSet dataset will be made publicly available on Github and Kaggle upon manuscript acceptance. Full access can be requested via email at during the review process.
